# *YES1* amplification: a mechanism of acquired resistance to EGFR inhibitors identified by transposon mutagenesis and clinical genomics

**DOI:** 10.1101/275974

**Authors:** Pang-Dian Fan, Giuseppe Narzisi, Anitha D. Jayaprakash, Elisa Venturini, Nicolas Robine, Peter Smibert, Soren Germer, Helena A. Yu, Emmet J. Jordan, Paul K. Paik, Yelena Y. Janjigian, Jamie E. Chaft, Lu Wang, Achim A. Jungbluth, Sumit Middha, Lee Spraggon, Huan Qiao, Christine M. Lovly, Mark G. Kris, Gregory J. Riely, Katerina Politi, Harold Varmus, Marc Ladanyi

**Affiliations:** Department of Pathology, Memorial Sloan Kettering Cancer Center, New York, NY 10065; Human Oncology and Pathogenesis Program, Memorial Sloan Kettering Cancer Center, New York, NY 10065; New York Genome Center, New York, NY 10013; Division of Solid Tumor Oncology, Department of Medicine, Memorial Sloan Kettering Cancer Center, New York, NY 10065; Vanderbilt-Ingram Cancer Center, Vanderbilt University School of Medicine, Nashville, TN 37232; Department of Pathology and the Yale Cancer Center, Yale University School of Medicine, New Haven, CT 06520; Cancer Biology and Genetics Program, Sloan Kettering Institute, Memorial Sloan Kettering Cancer Center, New York, NY 10065; Girihlet Inc., Oakland, CA 94609; University Hospital Waterford, Dunmore Road, Waterford, X91 ER8E, Ireland; Department of Pathology, St. Jude Children’s Research Hospital, Memphis, TN 38105; Meyer Cancer Center, Weill Cornell Medicine, New York, NY 10065

**Keywords:** YES1, EGFR, ALK, acquired resistance, lung adenocarcinoma, amplification, transposon mutagenesis, TKI

## Abstract

In approximately 30% of patients with *EGFR*-mutant lung adenocarcinomas whose disease progresses on EGFR inhibitors, the basis for acquired resistance remains unclear. We have integrated transposon mutagenesis screening in an *EGFR*-mutant cell line and clinical genomic sequencing in cases of acquired resistance to identify novel mechanisms of resistance to EGFR inhibitors. The most prominent candidate genes identified by insertions in or near the genes during the screen were *MET*, a gene whose amplification is known to mediate resistance to EGFR inhibitors, and the gene encoding the Src family kinase YES1. Cell clones with transposon insertions that activated expression of *YES1* exhibited resistance to all three generations of EGFR inhibitors and sensitivity to pharmacologic and siRNA-mediated inhibition of *YES1*. Analysis of clinical genomic sequencing data from cases of acquired resistance to EGFR inhibitors revealed amplification of *YES1* in 5 cases, 4 of which lacked any other known mechanisms of resistance. Pre-inhibitor samples, available for 2 of the 5 patients, lacked *YES1* amplification. None of 136 post-inhibitor samples had detectable amplification of other Src family kinases (*SRC, FYN*). *YES1* amplification was also found in 2 of 17 samples from *ALK* fusion-positive lung cancer patients who had progressed on ALK TKIs. Taken together, our findings identify acquired amplification of *YES1* as a novel, recurrent, and targetable mechanism of resistance to EGFR inhibition in *EGFR*-mutant lung cancers, and demonstrate the utility of transposon mutagenesis in discovering clinically relevant mechanisms of drug resistance.

**SIGNIFICANCE:** Despite high response rates to treatment with small molecule inhibitors of EGFR tyrosine kinase activity, patients with *EGFR*-mutant lung adenocarcinomas eventually develop resistance to these drugs. In many cases, the basis of acquired resistance remains unclear. We have used a transposon mutagenesis screen in an *EGFR*-mutant cell line and clinical genomic sequencing in cases of acquired resistance to identify amplification of *YES1* as a novel and targetable mechanism of resistance to EGFR inhibitors in *EGFR*-mutant lung cancers.

## INTRODUCTION

Four small molecule tyrosine kinase inhibitors (TKIs) have been FDA-approved for the treatment of *EGFR*-mutant lung cancers and represent three generations of drug development for this disease: erlotinib and gefitinib (1^st^ generation), afatinib (2^nd^) and osimertinib (3^rd^). Despite high response rates to these agents, the development of acquired resistance almost universally ensues. The mechanisms of acquired resistance can be grouped into target-dependent and target-independent categories. Target-dependent mechanisms are secondary alterations of *EGFR* that typically affect drug binding by, for example, altering the affinity of the kinase for ATP or by eliminating key sites for covalent bonding between drug and target protein. These include the T790M mutation that confers resistance to 1^st^ and 2^nd^ generation EGFR TKIs(1-4) and the C797S mutation that emerges upon osimertinib treatment(5, 6). Common target-independent mechanisms include amplification of *MET* and *ERBB2* (*HER2*) as well as small cell transformation(7, 8). However, in approximately 30% of cases of acquired resistance to 1^st^ generation EGFR TKIs, the underlying mechanisms still remain to be identified. While target-independent resistance mechanisms are expected to largely overlap between EGFR TKI generations, comprehensive studies of mechanisms of acquired resistance to 3^rd^ generation TKIs are currently ongoing.

To complement clinical genomic sequencing as a means of identifying mediators of resistance to EGFR inhibition, several different strategies have been employed using cell culture-based systems. Gradual escalation of concentrations of EGFR TKIs applied to *EGFR*-mutant lung cancer cell lines initially sensitive to the drugs has yielded TKI-resistant cells with clinically relevant mechanisms of resistance, including amplification of *MET*(9), overexpression of AXL(10), and secondary mutations of *EGFR*, most notably the T790M mutation(11-13). Forward genetic screens for modifiers of responses to EGFR inhibition, using libraries for RNA interference(14-18), expression of ORFs(16, 19), or CRISPR/Cas9-mediated gene deletion(16, 20), have also identified candidate genes that are implicated in acquired resistance in patients, including *NF1, BRAF, AXL*, and *ERBB2*.

Transposon-based mutagenesis is another forward genetic approach that can identify mechanisms of drug resistance. This strategy introduces genome-wide insertions of transposons, which have been designed with the potential to induce both gains and losses of endogenous gene function through the action of promoter/enhancer elements and splice acceptor and donor sequences that have been introduced into the transposons(21). Transposon mutagenesis has been used in cell culture-based systems and mouse models to screen for resistance to standard and investigational therapies for a variety of cancers, including paclitaxel(22), fludarabine(23), the PARP inhibitor olaparib(24), the MDM2-TP53 inhibitor HDM201(25), and the BRAF inhibitors PLX4720 and PLX4032(26, 27).

Here we report the results of an integrated approach, employing both forward genetic screening with transposon mutagenesis to recover drug-resistant derivatives of an *EGFR*-mutant lung adenocarcinoma cell line and genomic sequencing data from patients with acquired resistance to define novel, clinically relevant mechanisms of resistance to EGFR inhibition.

## RESULTS

### A transposon mutagenesis screen for resistance to EGFR inhibition in an *EGFR*-mutant lung adenocarcinoma cell line

To identify novel mechanisms of resistance to EGFR inhibition, we performed a transposon mutagenesis screen for resistance to the 2^nd^ generation EGFR TKI afatinib in the EGFR TKI-sensitive PC9 lung adenocarcinoma cell line, which harbors an activating small in-frame deletion in exon 19 of *EGFR* (**Fig. 1A**). Since transposon mutagenesis does not generate point mutations, our screen favored the recovery of target-independent mechanisms of resistance over target-dependent mechanisms such as the T790M and C797S second site mutations in *EGFR*. Although the emergence of some target-independent mechanisms of resistance might be suppressed by off-target TKI inhibition of kinases other than EGFR, we expected several of these mechanisms, including amplification of *MET*, to emerge repeatedly with successive generations of EGFR TKIs.

**Fig. 1.**
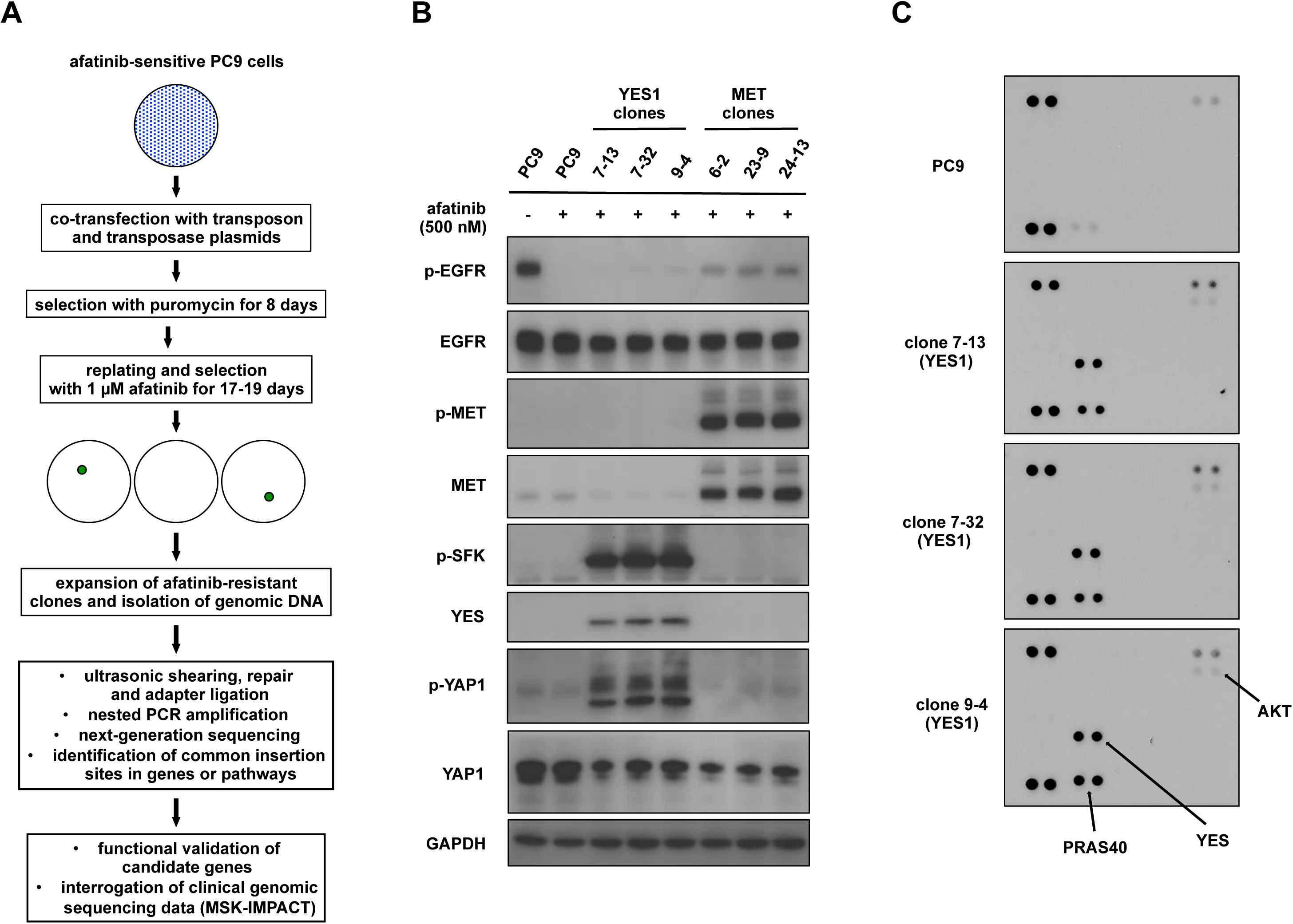
A transposon mutagenesis screen in *EGFR*-mutant PC9 lung adenocarcinoma cells for resistance to afatinib. (A) Flowchart representing the overall design of the screen. (B) Lysates from PC9 cells, *YES1* clones and *MET* clones treated with or without 500 nM afatinib for 60 minutes were subjected to immunoblot analysis with antibodies against the indicated proteins. (C) Lysates from PC9 cells and *YES1* clones treated with 500 nM afatinib for 60 minutes were hybridized to human phospho-kinase antibody arrays (R&D Systems, ARY003B).

PC9 cells were co-transfected with plasmids encoding a hyperactive piggyBac transposase(28) and a mutagenic transposon, which includes cytomegalovirus (CMV) enhancer and promoter sequences, a splice donor sequence, and a puromycin resistance cassette that provides a selection marker for transposon tagging(22). After selection with puromycin, transposon-tagged cells from 13 independent co-transfections were selected with 1 µM afatinib for 17 to 19 days. Afatinib-resistant clones were isolated for expansion and preparation of genomic DNA. No resistant clones were observed with non-transposon-tagged parental PC9 cells that were treated in parallel with 1 µM afatinib.

Transposon insertion sites were identified using a modified TraDIS-type method to generate Illumina-compatible libraries from DNA fragments that span the *piggyBac* sequence and the surrounding genomic DNA(29). Utilizing a custom bioinformatic pipeline with a set of filters based on the number of supporting reads, mean fragment size, and standard deviation of fragment size, we generated a list of 1927 distinct transposon insertion sites from 188 afatinib-resistant clones. Insertions were predicted to be activating if a transposon was situated near the transcription start site or first intron of a known human gene and was correctly oriented to drive expression of that gene. Genes that were found to be disrupted by insertions in both orientations or throughout the body of the gene were predicted to be inactivated.

### *MET* and *YES1* are the top candidate genes from the transposon mutagenesis screen for resistance to EGFR inhibition

Since the period between transfection and selection with afatinib was sufficient to allow one or more rounds of cell division of transposon-tagged cells, several clones from each transfection exhibited identical insertion sites, consistent with derivation from a common transfected progenitor. In selecting candidate genes for functional analysis, we therefore prioritized them based on the number of different insertions per gene and the number of independent transfections in which these insertions were discovered. The most promising candidate genes are listed in **Table 1**. The top two candidates were *MET*, encoding a receptor tyrosine kinase that is a known mediator of resistance, and *YES1*^*1*^, encoding a Src family kinase (SFK). All but one of the 188 clones harbored insertions in *MET* (78 clones), *YES1* (58 clones), or both genes (51 clones). In 29 clones, insertions were only found in *MET* out of the candidate genes listed in **Table 1**, and 45 clones had insertions in only *YES1* among these same candidate genes. The one clone that lacked insertions in either *MET* or *YES1* instead had insertions predicted to be activating in *SOS1* and *RABGAP1L*. Mutations in *SOS1* were recently found to be significantly enriched in lung adenocarcinoma samples without known driver alterations(30). As expected, *ERBB2*, another gene whose amplification is known to mediate resistance to erlotinib(31), was absent from the candidate list, reflecting the fact that afatinib also inhibits ERBB2(32).

**Table 1.** Candidate genes from a transposon mutagenesis screen for resistance to afatinib in the *EGFR*-mutant PC9 lung adenocarcinoma cell line. A total of 1927 distinct transposon insertion sites were identified in 188 afatinib-resistant PC9 clones from 13 independent transfections. Insertions were predicted to be activating if a transposon was situated near the transcription start site or first intron of a known human gene and was correctly oriented to drive expression of that gene. Genes that were found to be disrupted by insertions in both orientations or throughout the body of the gene were predicted to be inactivated.

| **Gene name** | **Predicted functional effect of insertions** | **Number of distinct insertion sites** | **Number of independent transfections with insertions** | **Total number of clones with insertions** |
| --- | --- | --- | --- | --- |
| *MET* | activating | 19 | 12 | 129 |
| *YES1* | activating | 15 | 8 | 109 |
| *FYN* | activating | 14 | 9 | 30 |
| *GSK3B* | inactivating | 7 | 4 | 8 |
| *SOS1* | activating | 6 | 12 | 37 |
| *DNM3* | inactivating | 4 | 8 | 24 |
| *NAV2* | inactivating | 4 | 5 | 15 |
| *EPS8* | activating | 4 | 2 | 6 |
| *KRAS* | activating | 4 | 2 | 5 |
| *MYOF* | inactivating | 3 | 4 | 19 |
| *RABGAP1L* | activating | 3 | 3 | 15 |
| *RAF1* | activating | 3 | 3 | 6 |
| *RABGAP1* | activating | 3 | 3 | 4 |

### Transposon insertions in *YES1* result in high expression and phosphorylation of YES1

We selected three clones with activating insertions in *MET* and another three with insertions in *YES1* - hereafter referred to as *MET* clones and *YES1* clones - for further characterization alongside parental PC9 cells. All six clones were maintained in growth medium containing 500 nM afatinib and lacked insertions in the other candidate genes listed in **Table 1**. To determine the levels of MET and YES1 proteins and phosphorylation of those proteins, we performed a series of immunoblots on cell lysates (**Fig. 1B**). High levels of phosphorylated MET were detected in *MET* clones. *YES1* clones exhibited high levels of YES1, phosphorylated SFKs, and phosphorylated Yes-associated protein 1 (YAP1). Since the phospho-SFK antibody does not distinguish between different SFKs, we analyzed cell lysates from *YES1* clones using a phospho-kinase array that specifically measures phosphorylation of YES, SRC, FYN and four other SFKs (**Fig. 1C**). In all three *YES1* clones, only phosphorylation of YES1 was detected among these seven SFKs. A similar survey using receptor tyrosine kinase (RTK) arrays showed phosphorylation of MET and ERBB3 in *MET* clones, and phosphorylation of ERBB3 in *YES1* clones, which was confirmed by immunoblot analysis (***SI Appendix*, Fig. S1**). Taken together, these findings confirm that the transposon insertions in *YES1* and *MET* resulted in high levels of the corresponding proteins; phosphorylation of these two kinases and their associated proteins is consistent with activation of YES1 and MET kinases in their respective clones.

### Clones with activating insertions in *YES1* are resistant to all three generations of EGFR TKIs, but are resensitized upon inhibition of YES1

We next determined if the *YES1* and *MET* clones were resistant to all three generations of EGFR inhibitors, and if the resistance was dependent on functional activity of YES1 and MET, respectively. Since only the 2^nd^ generation EGFR inhibitor afatinib was used in the transposon mutagenesis discovery screen, we tested the sensitivity of the clones to the 1^st^ generation TKI erlotinib and the 3^rd^ generation TKI osimertinib. Cell viability assays showed that all six clones were resistant to all three generations of EGFR inhibitors (**Fig. 2A and *SI Appendix*, Fig. S2A**). To block the kinase activities of YES1 and MET, we used the SFK inhibitors dasatinib and saracatinib and the MET inhibitor crizotinib, respectively. *YES1* clones were sensitive to the addition of dasatinib or saracatinib to afatinib, but not to the combination of crizotinib with afatinib (**Fig. 2B**). Conversely, *MET* clones were sensitive to the addition of crizotinib to afatinib, but not to the pairing of dasatinib or saracatinib with afatinib (***SI Appendix*, Fig. S2B**). *YES1* clones were also sensitive to the combination of either SFK inhibitor with osimertinib (**Fig. 2C**). Phosphorylation of serine-threonine kinase AKT and extracellular signal–regulated kinases (ERK) was observed in both *YES1* and *MET* clones, and was blocked by inhibiting SFKs or MET in addition to EGFR (**Fig. 2D**). Modest phosphorylation of EGFR, likely caused by kinases other than EGFR, was also abrogated by the addition of the SFK and MET inhibitors. Removal of afatinib from the growth medium for 72 hours restored high levels of phosphorylated EGFR in *YES1* clones, indicating that the intrinsic kinase activity of EGFR remained intact in these clones (***SI Appendix*, Fig. S2C**).

**Fig. 2.**
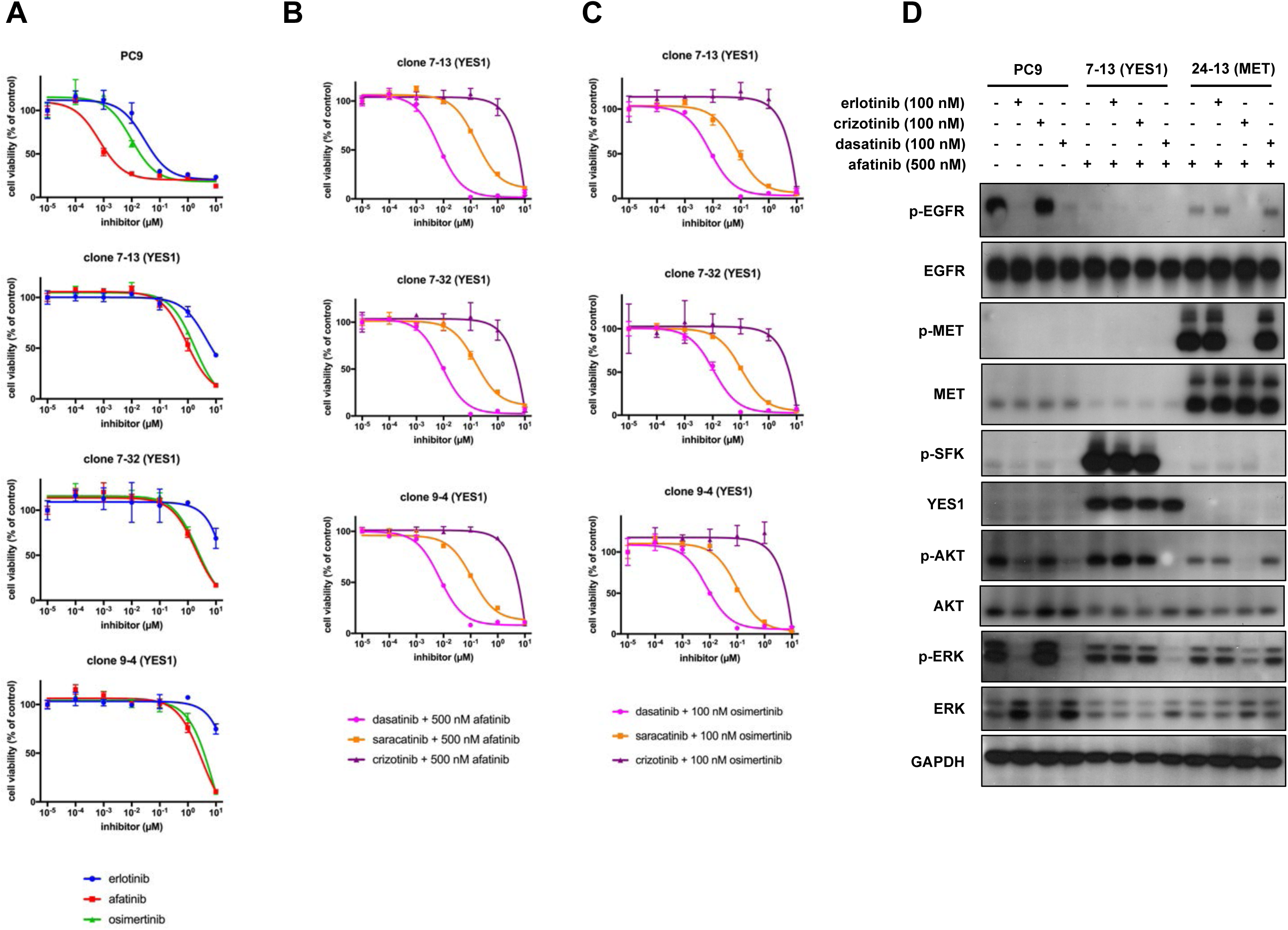
*YES1* clones are resistant to EGFR inhibitors from all three generations but sensitive when YES1 is inhibited. (A), (B), and (C) PC9 cells and *YES1* clones were seeded in 96-well plates and treated with EGFR inhibitors or the indicated inhibitors in combination with 500 nM afatinib or 100 nM osimertinib for 96 hours. Cell viability was assayed as described in Materials and Methods. Data are expressed as a percentage of the value for cells treated with a vehicle control and are means of triplicates. The experiments were performed 3 times with similar results. (D) Lysates from PC9 cells, clone 7-13 (*YES1*), and clone 24-13 (*MET*) treated with the indicated inhibitors for 60 minutes were subjected to immunoblot analysis with antibodies against the indicated proteins.

Since dasatinib and saracatinib have activity against kinases other than YES1, we specifically reduced YES1 levels by siRNA-mediated knockdown and assessed the effect on the viability of *YES1* clones. In addition, since the FLAURA study recently showed superior efficacy of osimertinib to that of standard EGFR TKIs in the first-line treatment of EGFR mutation-positive advanced NSCLC, we chose osimertinib to combine with siRNA-mediated knockdown of *YES1*(33). As shown in **Fig. 3A**, the *YES1*-specific siRNA, but not the negative control or *MET*-specific siRNA, sensitized *YES1* clones to treatment with osimertinib. In contrast, the *MET*-specific siRNA, but not the negative control or *YES1*-specific siRNA, sensitized *MET* clones to treatment with osimertinib (**Fig. 3B**). Neither the *YES1*-specific siRNA or *MET*-specific siRNA increased the sensitivity of parental PC9 cells to treatment with osimertinib (**Fig. 3C**). These results are consistent with YES1 as the key target of SFK inhibitors in *YES1* clones and confirm that YES1 is required to mediate the resistance of these clones to EGFR inhibitors.

**Fig. 3.**
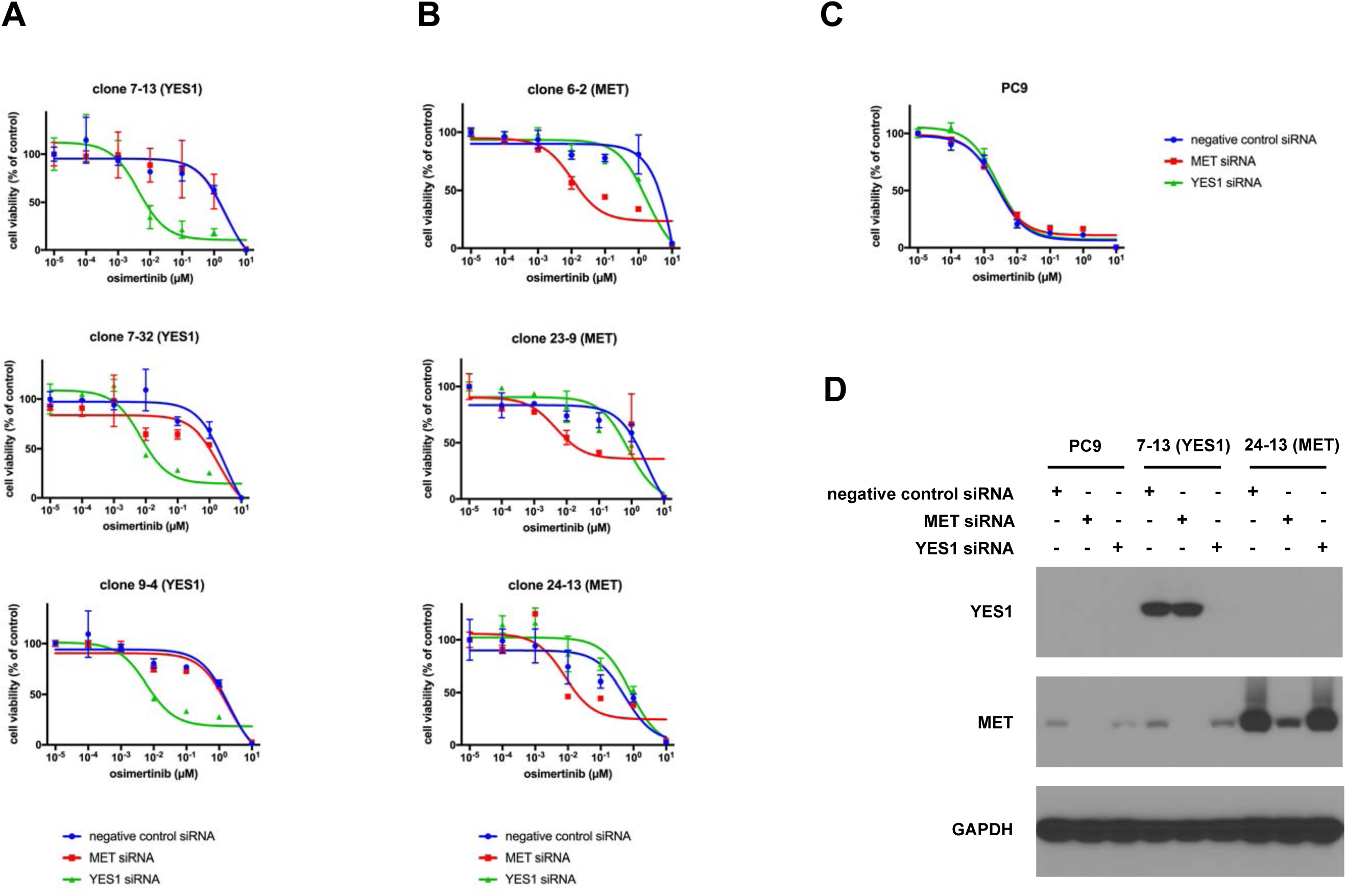
*YES1* clones are resistant to osimertinib but are resensitized by siRNA-mediated knockdown of *YES1*. *YES1* clones (A), *MET* clones (B), and PC9 cells (C) were transfected with negative control, *MET*-specific, and *YES1*–specific siRNAs at a final concentration of 10 nM. After 24 hours, cells were trypsinized and seeded in 96-well plates at a density of 5,000 cells per well with the indicated concentrations of osimertinib for 72 hours followed by measurement of cell viability. Experiments were performed 3 times with similar results. (D) Immunoblot analysis with YES1, MET, and GAPDH antibodies was performed on lysates prepared from PC9 cells, clone 7-13, and clone 24-13 72 hours after transfection with the indicated siRNAs.

### *YES1* is amplified in clinical cases of acquired resistance to EGFR inhibitors

To search for clinical evidence of a role for *YES1* in acquired resistance to EGFR inhibition, we examined clinical genomic sequencing data generated with the MSK-IMPACT panel from 136 patients whose *EGFR*-mutant lung adenocarcinomas progressed on EGFR inhibitors(34). This data set included 128 post-erlotinib, 6 post-afatinib, and 2 post-dacomitinib cases of acquired resistance to EGFR inhibition. Amplification of *YES1* was identified in 3 out of 66 T790M-negative cases and 1 out of 70 T790M-positive cases. None of the 136 cases had detectable amplification of *SRC* or *FYN*, the two other SFKs included in the MSK-IMPACT panel assay. The MSK-IMPACT fold changes (normalized log_2_ transformed fold changes of coverage of tumor versus normal) and FACETS integer values (allele-specific copy numbers corrected for tumor purity, ploidy and clonal heterogeneity) for *YES1* are listed in **Table 2**, and all the copy number profiles are shown in **Fig. 4A** and ***SI Appendix*, Fig. S3**. The responses to the indicated EGFR TKIs in the 4 cases with amplification of *YES1* ranged from approximately 5 months (patient 1) to approximately 30 months (patient 2). Additionally, separate from this cohort of 136 consecutive patients with MSK-IMPACT data on their *EGFR*-mutant lung adenocarcinomas with acquired resistance to EGFR inhibitors, we recently detected a striking level of acquired *YES1* amplification in a fifth *EGFR*-mutant case, a T790M-negative case of progression of disease on erlotinib and maintenance pemetrexed. Previous treatment regimens in this patient also included the use of carboplatin, bevacizumab, and localized radiation therapy to the site of progression. The progression sample also harbored a missense mutation of unclear significance in *YES1*, Q322H, that appeared to be amplified copies of the gene based on its variant allele fraction (patient 5, **Table 2** and **Fig. 4A**). A prior post-TKI sample two years earlier did not show *YES1* amplification.

**Table 2.**
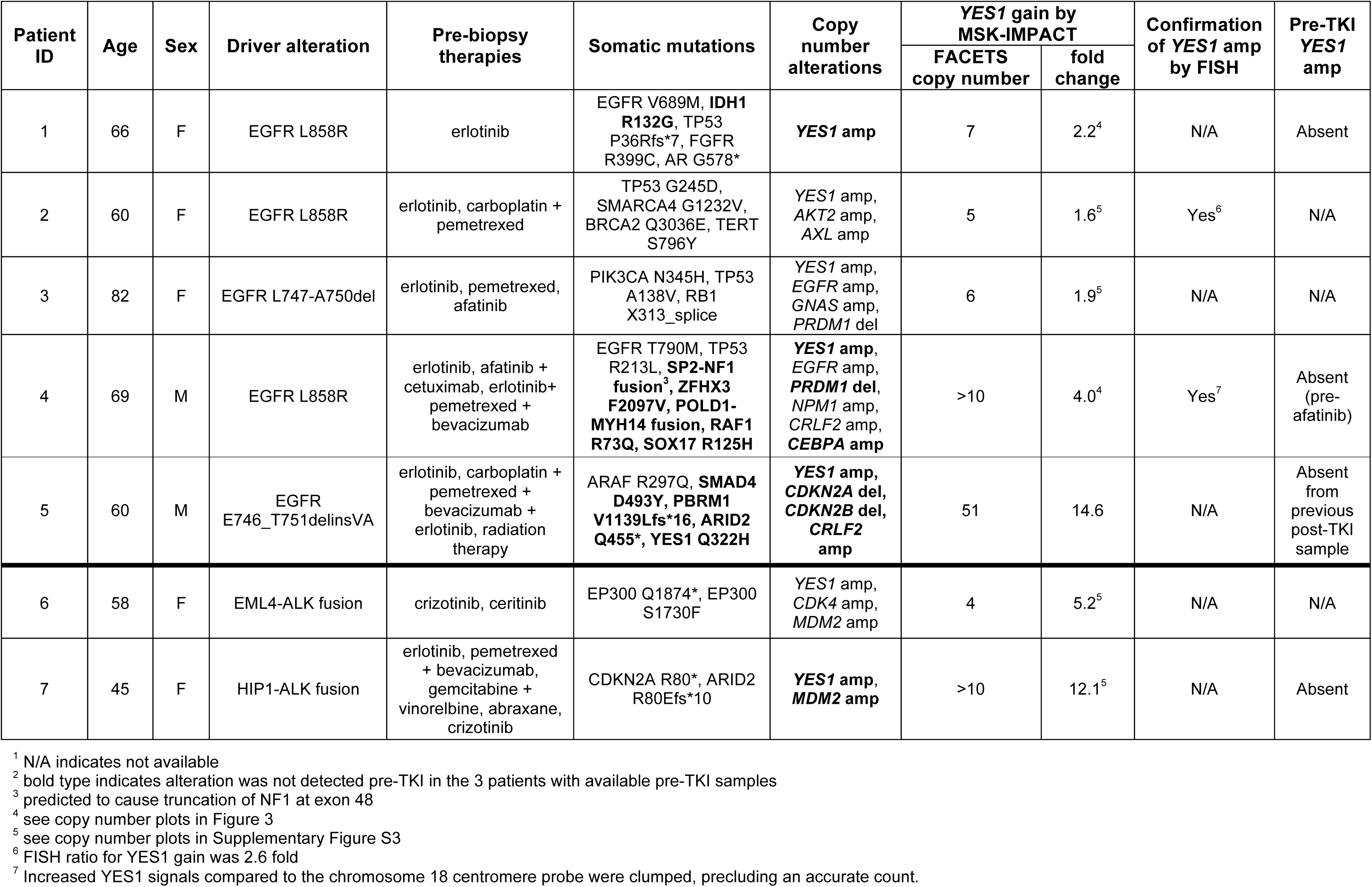
Clinical and molecular features of cases of acquired resistance to EGFR or ALK inhibitors with amplification of *YES1*. The FACETS integer values are allele-specific copy numbers corrected for tumor purity, ploidy and clonal heterogeneity. The MSK-IMPACT fold changes are normalized log_2_ transformed fold changes of coverage of tumor versus normal.

**Fig. 4.**
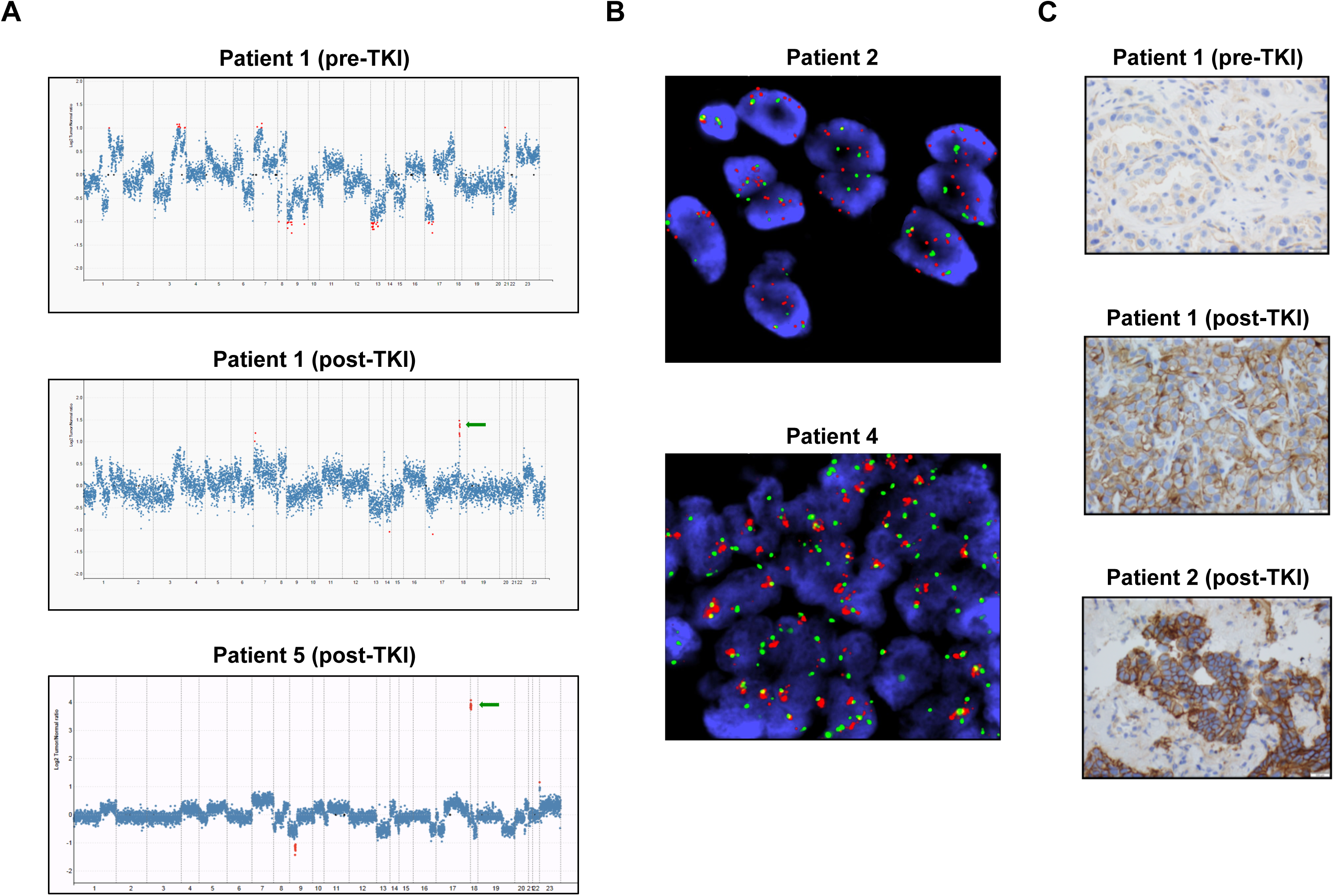
Amplification of *YES1* in tumor samples from patients with acquired resistance to EGFR inhibitors. (A) Copy number plots for tumor samples from patients 1 and 5. Each dot represents a target region in the MSK-IMPACT targeted capture assay. Red dots are target regions exceeding a fold change cutoff of 2-fold. The log-ratios (y-axis) comparing tumor versus normal coverage values are calculated across all targeted regions (x-axis). Green arrows indicate focal amplification of *YES1* (11 coding exons targeted). (B) *YES1* FISH for post-TKI tumor specimens from patients 2 and 4. *YES1* (red) and *CEP18* (green). For patient 2, the FISH ratio for *YES1* gain was 2.6 fold. For patient 4, increased *YES1* signals were clumped, precluding an accurate count. (C) Immunohistochemistry for YES1 on tumor samples from patients 1 and 2. The clinical and molecular features of these patients are summarized in Table 2.

Although the MSK-IMPACT assay is not designed to enable a formal analysis of minimal regions of gain or loss, additional focality data were available based on the assay results. *YES1* is the most telomeric gene on 18p included in the MSK-IMPACT assay, extending from position 724,421 to 756,830. In all 7 cases in Table 2, all *YES1* exons showed an increase in copy number. The next set of probes immediately centromeric to *YES1* are intergenic tiling probes extending from 2,224,682 to 38,530,030. The next closest gene in the assay panel is *PIK3C3* starting at 39,535,254; none of the 7 cases showed co-amplification of *PIK3C3*. In 2 of the 7 cases in Table 2 (patients 2 and 3), the *YES1* gains included some of the tiling probes on the centromeric side, with the furthest being 34,882,991 in the former case. In the remaining 5 cases, amplification was only detected with the *YES1* probes.

Amplification of *YES1* was confirmed by fluorescence *in situ* hybridization (FISH) for two cases with sufficient material for analysis (**Fig. 4B**). Immunohistochemical staining for YES1 in post-TKI samples from patients 1 and 2 showed prominent labeling of lung adenocarcinoma cells which, moreover, was absent in the pre-TKI specimen available from patient 1 (**Fig. 4C**). A previously known mechanism of resistance was found in only one out of the four samples containing amplified *YES1*, namely the EGFR T790M mutation, but with *EGFR* mutation allele frequencies (L858R 0.77, T790M 0.16) that were consistent with intratumoral heterogeneity, raising the possibility that the T790M *EGFR* allele and the amplified *YES1* allele were in separate subpopulations, as has been described in other instances of multiple concurrent resistance mechanisms(7, 8, 35, 36). In addition, amplification of *YES1* was not detected in pre-TKI samples that were available for patients 1 and 4, confirming that it had emerged during treatment. Interestingly, review of the MSK-IMPACT data in other molecular subsets of lung adenocarcinoma revealed *YES1* amplification in 2 out of 17 ALK fusion-driven lung adenocarcinomas that had acquired resistance to ALK TKIs. These two cases did not show evidence of other changes, such as secondary mutations in *ALK* that have previously been found in such tumors that developed resistance to ALK inhibitors (**Table 2**). In one of these cases, the pre-TKI sample was available and showed no *YES1* amplification.

To assess the occurrence of *YES1* amplification generally in lung adenocarcinomas, not just those with acquired resistance to EGFR and ALK inhibitors, we reviewed all 2466 lung adenocarcinomas in a more recent version of the MSK-IMPACT patient database (data freeze: August 31, 2017). In addition to the previously described 4 *EGFR*-mutant and 2 *ALK*-rearranged lung adenocarcinomas, we found 13 more cases with an amplified *YES1* locus, including 2 *EGFR*-mutant tumors pre-TKI treatment, 3 tumors with a *KRAS* mutation, 1 pre-TKI tumor with a *MET* mutation causing exon 14 skipping, and 8 tumors without a known driver mutation. These data indicate that *YES1* amplification is rarely detected prior to targeted therapy for *EGFR*-mutant and *ALK*-rearranged lung adenocarcinomas, and is not commonly found in lung adenocarcinomas in general.

## DISCUSSION

The present approach of integrating transposon mutagenesis screening data in lung adenocarcinoma cell lines with clinical genomic sequencing data from patient tumor specimens identified both established and novel mechanisms of resistance to EGFR inhibition. The most prominent candidate genes from the screen for resistance to afatinib in PC9 cells were *MET* and *YES1*. While our screen with afatinib was initiated well before the FDA approval of the 3^rd^ generation EGFR TKI osimertinib, it is important to note that clones with activating transposon insertions in these genes were resistant to erlotinib, afatinib, and osimertinib, representing respectively all three generations of FDA-approved EGFR TKIs.

Review of the MSK-IMPACT patient database revealed post-TKI amplification of *YES1* in 5 *EGFR*-mutant and 2 *ALK*-rearranged lung adenocarcinomas with acquired resistance to targeted therapy. Only one of these 7 post-TKI samples harbored another known resistance-conferring alteration, specifically a concurrent EGFR T790M mutation. The presence of a concurrent EGFR T790M in one case is not unexpected, as biopsy samples in this clinical setting can show more than one resistance mechanism, presumably due to intratumoral heterogeneity. Amplification of *YES1* was not detected in pre-TKI samples that were available for two of the *EGFR*-mutant cases and one of the *ALK*-rearranged cases, confirming its acquired nature. To our knowledge, this is the first report of genomic sequencing data from patient tumors that implicate amplification of SFK genes in therapeutic resistance.

Previous laboratory studies support amplification of *YES1* as a mediator of resistance to inhibitors of ERBB family members. Ichihara and colleagues found that amplification of *YES1* mediated resistance in 1 out of 5 PC9-derived cell lines that were rendered resistant in culture to osimertinib through gradual dose escalation(18). Amplification of *YES1* has also been shown to mediate resistance to trastuzumab and lapatinib in drug-resistant models that were generated from an *ERBB2*-amplified breast cancer cell line(37). In addition, an initial ORF-based screen for genetic modifiers of EGFR dependence in PC9 cells identified 8 of the 9 SFK genes as potentially reducing sensitivity to erlotinib(19). *YES1* was the only one of the 8 SFK genes to fail subsequent validation assays, but its functional validation may have been hampered by the markedly lower level of YES1 protein achievable experimentally in comparison to the other 7 SFKs. Similarly, despite utilizing multiple vectors featuring either constitutive or tetracycline-regulated promoters, we have also been unable to achieve robust ectopic expression of YES1 in PC9 cells. Given these technical limitations, further functional studies in PC9 cells will likely require approaches to upregulate YES1 expression from its endogenous locus.

Treatment with SFK inhibitors, such as dasatinib, has previously been investigated in the setting of acquired resistance to EGFR TKIs. Johnson and colleagues did not observe activity for the combination of erlotinib and dasatinib in 12 patients with *EGFR*-mutant lung adenocarcinoma with acquired resistance to erlotinib or gefitinib, of which 8 were positive for T790M prior to initiation of this combination therapy(38). However, the copy number status of *YES1* was not determined in this trial and the likelihood that one of the four T790M-negative patients in this trial had *YES1* amplification as a resistance mechanism is statistically very low. Our findings justify consideration of treatment with combined EGFR and SFK inhibition in the subset of cases of acquired resistance to EGFR TKIs that harbor amplification of *YES1*. Our results also suggest that this mechanism might contribute to resistance to ALK TKIs. In this context, it is notable that a pharmacogenomic screen of cell lines derived from ALK TKI-resistant *ALK* fusion-positive lung cancer biopsies recently identified several cell lines that exhibited upregulated SRC activity upon ALK inhibition and sensitivity to dual ALK and SRC inhibition(39). Although the mechanism of SRC regulation by ALK signaling remains unclear, these data suggest an important role for signaling downstream of SFKs in a subset of ALK TKI-resistant *ALK* fusion-positive lung cancers.

We have shown that transposon mutagenesis screening can facilitate identification of clinically relevant target-independent mechanisms of resistance to EGFR inhibition. This approach can be rapidly re-implemented to screen *in vitro* for resistance to additional drugs or drug combinations. By incorporating additional EGFR TKIs (e.g., osimertinib), other *EGFR*-mutant cell lines, and concurrent inhibition of EGFR, MET and YES1 in PC9 cells, reapplication of transposon mutagenesis has the potential to clarify the contributions of other candidate genes identified in our screen to resistance to EGFR inhibition and to uncover additional mediators of resistance to EGFR inhibitors. We anticipate that other targeted therapies in lung adenocarcinomas will be amenable to this approach to identifying novel mechanisms of resistance.

## Materials and Methods

### Cell culture and inhibitors

PC9 cells were obtained from the Varmus laboratory and have been maintained in the Ladanyi laboratory since 2010. Cells were cultured in RPMI-1640 medium supplemented with 10% FBS (Atlanta Biologicals) and 100 U/mL penicillin/100 µg/mL streptomycin (Gemini Bio-Products). Cells were grown in a humidified incubator with 5% CO_2_ at 37°C. Afatinib-resistant clones were maintained in growth medium with 500 nM afatinib. Afatinib, erlotinib, osimertinib, dasatinib, saracatinib and crizotinib were obtained from Selleck Chemicals.

### Transposon mutagenesis

PC9 cells were seeded at a density of 5×10^5^ cells per well in 6-well plates 24 hours prior to co-transfection with plasmids pCMV-HA-hyPBase (obtained from the Wellcome Trust Sanger Institute) and pPB-SB-CMV-puroSD (obtained from Li Chen, Schmidt Laboratory, Massachusetts General Hospital) using X-tremeGENE™ 9 DNA transfection reagent (Roche) according to the manufacturer’s protocol. After 48 hours, cells were selected in growth medium with 0.3 µg/ml puromycin for 8 days. Surviving cells from 13 independent co-transfections were replated at a density of 2.7×10^5^ cells per 15-cm plate in growth medium with 1 µM afatinib for 17 to 19 days. A total of 225 resistant clones were isolated using cloning discs (Scienceware) and expanded. Genomic DNA was prepared from clones using DNeasy Blood & Tissue Kits (Qiagen).

## Library preparation, next-generation sequencing and bioinformatics analysis for identification of transposon insertion sites. See *SI Materials and Methods*

### Immunoblot and phospho-kinase array analyses

Cells were washed with ice-cold PBS and lysed with RIPA buffer (Cell Signaling Technology) supplemented with protease inhibitor and phosphatase inhibitor cocktails (Roche). Protein levels were quantified with Bradford dye reagent (Bio-Rad), and equal amounts were loaded for SDS-PAGE using precast Bis-Tris gels (Invitrogen), followed by transfer to polyvinylidene difluoride membranes. Membranes were blotted with the following antibodies according to the supplier’s recommendations and were all obtained from Cell Signaling Technology unless otherwise noted: phospho-EGFR Y1068 (#3777), EGFR E746-A750del specific (#2085), phospo-MET Y1234/1235 (#3777), MET (#8198), phospho-SFK (#6943), YES (#3201), phospho-YAP1 Y357 (#ab62751, Abcam), YAP1 (#14704), phospho-AKT S473 (#4060), AKT (#4691), phospho-ERK T202/Y204 (#4370), ERK (#4695), phospho-ERBB3 Y1197 (#4561), ERBB3 (#12708), GAPDH (#2118). Human phospho-kinase (#ARY003B, R&D Systems) and human phospho-RTK ((#ARY001B, R&D Systems) array kits were used according to the manufacturer’s protocols.

### Cell viability assays

Cells were seeded in 96-well plates at a density of 2,500 (PC9) to 5,000 (*YES1* and *MET* clones) cells per well in growth medium with the indicated inhibitors. After 96 hours, AlamarBlue cell viability reagent (Invitrogen) was added at a final concentration of 10% (vol/vol) and fluorescence was measured (Ex: 555 nm, Em: 585) with a SpectraMax M2 plate reader.

### siRNA experiments

Cells were transfected with negative control and *YES1*– specific siRNAs (Invitrogen) at a final concentration of 10 nM using Lipofectamine RNAiMAX reagent (Invitrogen) according to the manufacturer’s protocol. After 24 hours, cells were trypsinized and seeded in 96-well plates at a density of 5,000 cells per well and incubated with the indicated inhibitors for 72 hours followed by measurement of cell viability.

### Statistical analysis

Mean and standard deviation values for cell viability assays were calculated and plotted using Prism 7 software (GraphPad Software). Copy number aberrations were identified using an in-house developed algorithm by comparing sequence coverage of targeted regions in a tumor sample relative to a standard diploid normal sample(40), as extensively validated for *ERBB2* (*HER2*) amplification(41). Allele-specific copy number alterations were also identified using the FACETS analysis tool, which performs a joint segmentation of the copy ratios(42). For a complete list of genes included in the MSK-IMPACT panel, see Table A1 in the cited reference(43).

### FISH analysis

Interphase FISH analysis on formalin-fixed paraffin-embedded (FFPE) tumor tissue was performed to evaluate *YES1* gene copy number status. The probe targeting *YES1* at 18p11.32 was labeled with SpectrumOrange fluorochrome (Empire Genomic) and the control probe targeting the centromere of chromosome 18 was labeled with SpectrumGreen (Abbott Molecular). Four-micron (4 µm) FFPE tissue sections were used for the FISH study, following the protocol for FFPE tissue FISH from Vysis/Abbott Molecular with minor adjustments of pepsin treatment as needed. FISH analysis and signal capture were conducted on a fluorescence microscope (Zeiss) coupled with the ISIS FISH Imaging System (Metasystems). We analyzed 100 interphase nuclei from tumor-rich areas in each specimen.

### Immunohistochemistry

For the immunohistochemical detection of YES1, monoclonal antibody EPR3173 (Abcam, 1:250) was used. All staining procedures were performed on a Leica Bond-3 automated stainer platform. Heat-based antigen retrieval using a high pH buffer (ER2, Leica) was employed before the actual staining. A polymer-based secondary system (Leica Refine) was used to detect the primary antibody.

## Author contributions

P.F., K.P., H.V., and M.L. designed research. P.F., P.S., A.D.J., G.N., E.V., N.R., S.G., H.A.Y., E.J.J., P.K.P., Y.Y.J., J.E.C., L.W., A.A.J., S.M., L.S., H.Q., and G.J.R. performed research. P.F., P.S., A.D.J., G.N., N.R., S.M., C.M.L., M.G.K., and M.L. analyzed data. P.F. and M.L. wrote the paper with input from all authors.

## ACKNOWLEDGEMENTS

We thank Li Chen and Eiki Ichihara for reagents, Venkatraman Seshan for assistance with FACETS analysis, and Mary Ann Melnick for technical assistance. This project was begun as a collaboration with K.P. when P.F. was a postdoctoral fellow in the laboratory of H.V. at MSKCC, and was continued by P.F. in the laboratory of M.L. This work was supported by National Institutes of Health (NIH)/National Cancer Institute (NCI) grants R01 CA120247 (to H.V. and K.P.), P01 CA129243 (to M.G.K. and M.L.), and P30 CA008748 (MSKCC); the New York State Empire Clinical Research Investigator Program (P.F.); the Lung Cancer Research Foundation (P.F. and M.L.); the Functional Genomics Initiative at MSKCC (P.F., M.L., G.J.R. and L.W.); and the Katha Diddel Sussman & Warren Family Fund for Genome Research (P.F.). C.M.L.was additionally supported by a V Foundation Scholar-in-Training Award, an AACR-Genentech Career Development Award, a Damon Runyon Clinical Investigator Award, a LUNGevity Career Development Award, and NIH/NCI R01 CA121210.

## Supporting Information

### SI Materials and Methods

#### Library preparation and sequencing

*piggyBac* insertion sites were identified by generating Illumina-compatible libraries from DNA fragments that span the *piggyBac* sequence and the surrounding genomic DNA, using a modified TraDIS-type method(29). Genomic DNA (200 ng) from 188 isolated clones was sheared in a Covaris 96 microTUBE Plate to give fragments around 350 base pairs in length. End repair, A-tailing and ligation were performed using the Bioo NextFlex rapid DNA sequencing kit. Following ligation and removal of excess adapters, a two-step nested PCR protocol was performed to create sequencing libraries. For the first PCR (12 cycles), primers F-nest_1 (designed to bind the adapter sequence) and Pb-51 (designed to bind the *piggyBac* sequence) were used to specifically amplify fragments that contain the *piggyBac* end sequence. Products were purified, and re-amplified with F-X and N2-X primers (16 cycles) to append indices and library ends. Using a dual indexing strategy, the combination of 8 F-X primers and 12 N2-X primers allows 96 index combinations. The F-X primer binds to the adapter sequence, and adds the i5 index and the P5 end of the library. The N2-X primer binds to the *piggyBac* sequence at a position immediately adjacent and proximal to the binding site of the Pb-51 primer used in the first amplification. It also contains the i7 index sequence and the P7 sequence for binding to the Illumina flowcell. PCRs were performed with Phusion polymerase using high GC buffer. Sequencing was performed on a Miseq using a 2×75bp run with custom sequencing primers for index 1 and read 2. The PCR strategy was designed so that for read 2, the first 17 nucleotides sequenced (tatctttctagggttaa) correspond to the end of *piggyBac*, to enable unambiguous identification of *piggyBac* end sequences and precise delineation of insertion sites.

Primer sequences:

F-nest_1: TCTTTCCCTACACGACGCTCTTCCGATCT

Pb-51: CGCTATTTAGAAAGAGAGAGCAATATTTCA

F-X: AATGATACGGCGACCACCGAGATCTACACxxxxxxxxACACTCTTTCCCTACACGACGCTCTTCCGATCT

N2-X: CAAGCAGAAGACGGCATACGAGATxxxxxxAGAATGCATGCGTCAATTTTACGCAGAC

Custom index 1 sequencing primer: GTCTGCGTAAAATTGACGCATGCATTCT

Custom read 2 sequencing primer: AGAATGCATGCGTCAATTTTACGCAGAC

F-1: AATGATACGGCGACCACCGAGATCTACACTAGATCGCACACTCTTTCCCTACACGACGCTCTTCCGATCT

F-2: AATGATACGGCGACCACCGAGATCTACACCTCTCTATACACTCTTTCCCTACACGACGCTCTTCCGATCT

F-3: AATGATACGGCGACCACCGAGATCTACACTATCCTCTACACTCTTTCCCTACACGACGCTCTTCCGATCT

F-4: AATGATACGGCGACCACCGAGATCTACACAGAGTAGAACACTCTTTCCCTACACGACGCTCTTCCGATCT

F-5: AATGATACGGCGACCACCGAGATCTACACGTAAGGAGACACTCTTTCCCTACACGACGCTCTTCCGATCT

F-6: AATGATACGGCGACCACCGAGATCTACACACTGCATAACACTCTTTCCCTACACGACGCTCTTCCGATCT

F-7: AATGATACGGCGACCACCGAGATCTACACAAGGAGTAACACTCTTTCCCTACACGACGCTCTTCCGATCT

F-8: AATGATACGGCGACCACCGAGATCTACACCTAAGCCTACACTCTTTCCCTACACGACGCTCTTCCGATCT

N2-1: CAAGCAGAAGACGGCATACGAGATTGGTCAAGAATGCATGCGTCAATTTTACGCAGAC

N2-2: CAAGCAGAAGACGGCATACGAGATCACTGTAGAATGCATGCGTCAATTTTACGCAGAC

N2-3: CAAGCAGAAGACGGCATACGAGATCTGATCAGAATGCATGCGTCAATTTTACGCAGAC

N2-4: CAAGCAGAAGACGGCATACGAGATTACAAGAGAATGCATGCGTCAATTTTACGCAGAC

N2-5: CAAGCAGAAGACGGCATACGAGATCGTACGAGAATGCATGCGTCAATTTTACGCAGAC

N2-6: CAAGCAGAAGACGGCATACGAGATCCACTCAGAATGCATGCGTCAATTTTACGCAGAC

N2-7: CAAGCAGAAGACGGCATACGAGATATCAGTAGAATGCATGCGTCAATTTTACGCAGAC

N2-8: CAAGCAGAAGACGGCATACGAGATGCCTAAAGAATGCATGCGTCAATTTTACGCAGAC

N2-9: CAAGCAGAAGACGGCATACGAGATCGTGATAGAATGCATGCGTCAATTTTACGCAGAC

N2-10: CAAGCAGAAGACGGCATACGAGATACATCGAGAATGCATGCGTCAATTTTACGCAGAC

N2-11: CAAGCAGAAGACGGCATACGAGATATTGGCAGAATGCATGCGTCAATTTTACGCAGAC

N2-12: CAAGCAGAAGACGGCATACGAGATAAGCTAAGAATGCATGCGTCAATTTTACGCAGAC

index sequences (replacing “X” in generic primer sequences above)

N2-1: TGGTCA

N2-2: CACTGT

N2-3: CTGATC

N2-4: TACAAG

N2-5: CGTACG

N2-6: CCACTC

N2-7: ATCAGT

N2-8: GCCTAA

N2-9: CGTGAT

N2-10: ACATCG

N2-11: ATTGGC

N2-12: AAGCTA

F-1: TAGATCGC

F-2: CTCTCTAT

F-3: TATCCTCT

F-4: AGAGTAGA

F-5: GTAAGGAG

F-6: ACTGCATA

F-7: AAGGAGTA

F-8: CTAAGCCT

### Bioinformatics analysis

Illumina paired-end reads were aligned to the reference genome (hg19) using bwa mem v0.7.8 [1] with default parameters. PCR duplicates were marked using picard tools^2^ v1.83 and were not used for downstream analysis. In order to generate a list of candidate *piggyBac* insertion sites we developed custom software (using the BamTools API^3^) to analyze the pattern of soft-clipped sequences in the aligned reads. Reads were extracted if the soft-clipped portion of the sequence (either forward or reverse complemented) contained a perfect match (≥ 13 bp) to the *piggyBac* sequence. A candidate insertion site was considered if all soft-clipped sequences (matching *piggyBac*) for a predefined minimum number of reads start at the same coordinate in the genome. Read orientation and alignment position of the mate are collected for all reads supporting the insertion site and used to infer the orientation of the insertion and compute statistics to remove false-positive candidates. For actual *piggyBac* insertion sites one would expect the insert size distribution, and the standard deviation (std) of insert sizes, to follow the expected distribution, measured by mapping the read pairs to the reference genome. However, due to ambiguity in read mapping or library artifacts, this may not hold. **Fig. S4A** and **Fig. S4B** show that most of the candidate insertions sites follow the expected insert size distribution inferred after alignment (mean=100, std=55) and **Fig. S4C** shows good correlation between mean and std of the fragment insert size. The exception is a suspiciously high number of sites characterized by a small std. These sites are likely to be false-positives and in fact the correlation analysis (**Fig. S4C**) shows that most of them have significantly higher mean insert size. Based on these results we compiled a list of high confidence insertion sites using a conservative set of filters: >10 supporting reads (matching the *piggyBac* sequence) and insert size std > 10. After filtering, our list contained 1927 distinct candidate insertion sites in 188 clones. We then focused our attention to genes with multiple independent insertion events.

## SI Figure Legends

**Fig. S1.**
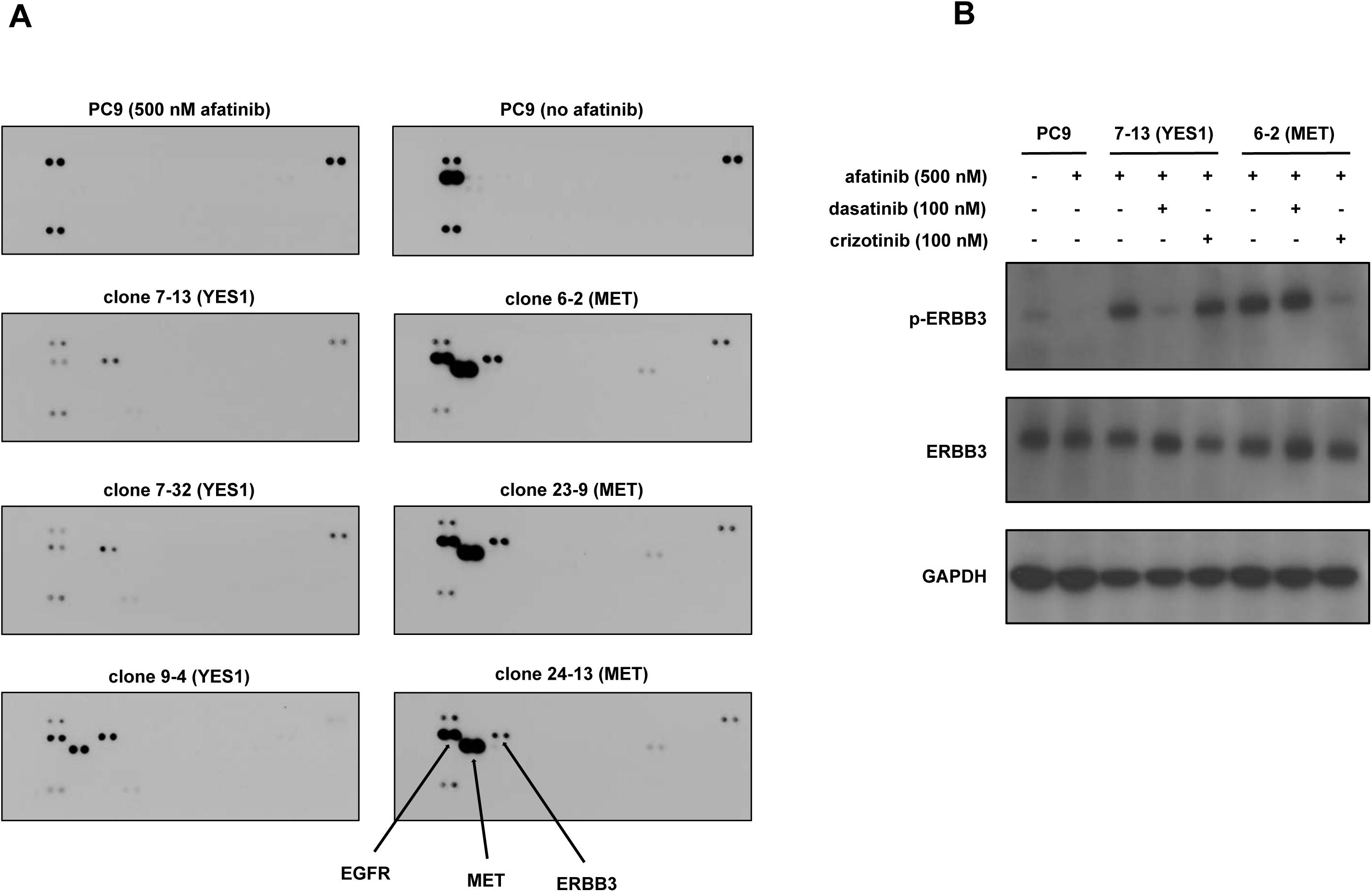
Tyrosine phosphorylation of ERBB3 in *YES1* and *MET* clones. (A) Lysates from PC9 cells with and without 500 nM afatinib and from *YES1* and *MET* clones maintained in 500 nM afatinib were hybridized to human phospho-RTK arrays (R&D Systems, ARY001B). The pan phospho-tyrosine antibody for the array kit detects phosphorylation of MET in clone 9-4, presumably at a site different from the tyrosine recognized by the phospho-MET antibody used in Figure 1B. (B) Lysates from PC9 cells, clone 7-13 (*YES1*), and clone 6-2 (*MET*) treated with the indicated inhibitors for 60 minutes were subjected to immunoblot analysis with antibodies against the indicated proteins.

**Fig. S2.**
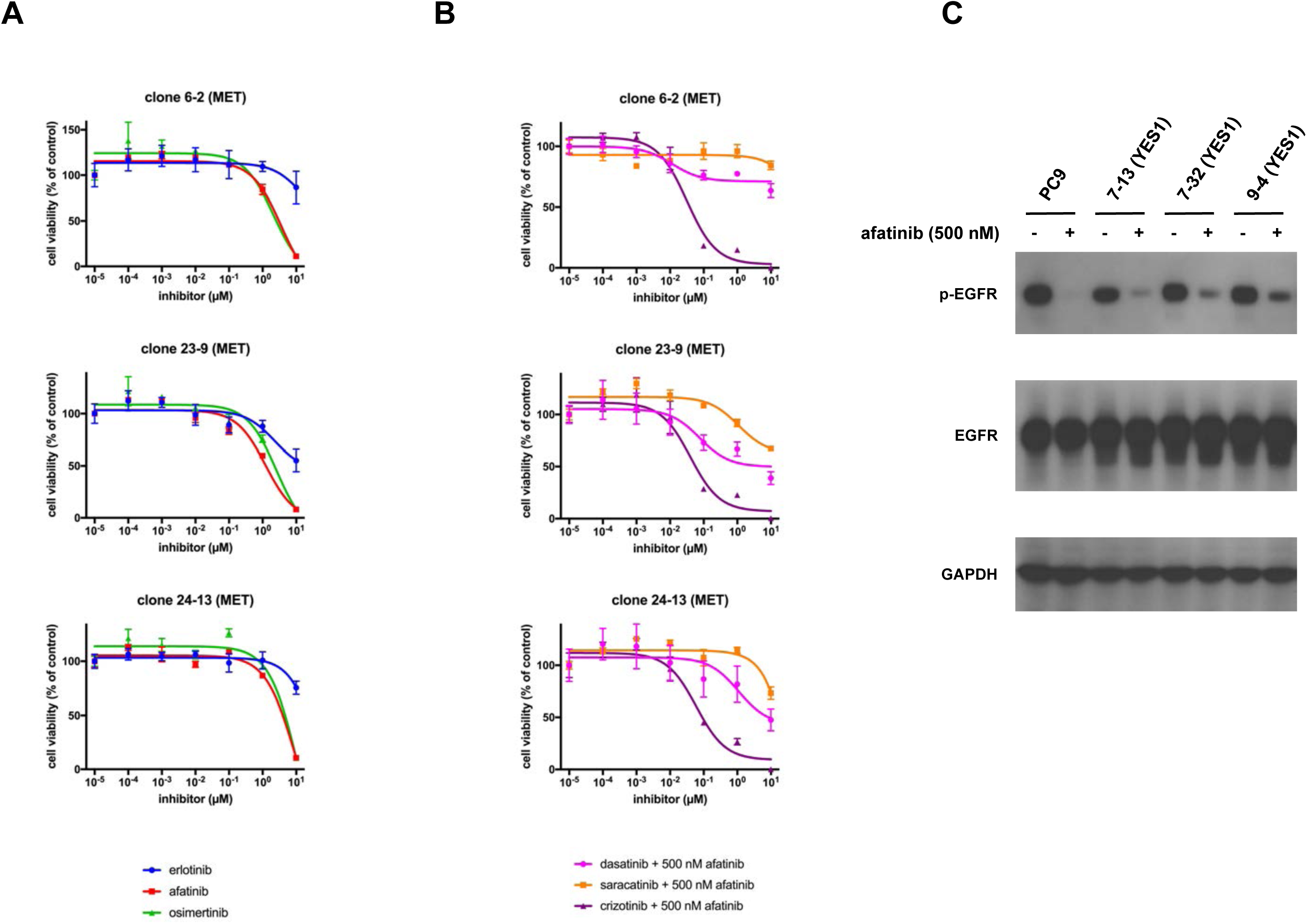
*MET* clones are resistant to EGFR inhibitors from all three generations but sensitive to additional blockade of MET kinase activity. *YES1* clones retain EGFR kinase activity. (A) and (B) *MET* clones were seeded in 96-well plates and treated with EGFR inhibitors or the indicated inhibitors in combination with 500 nM afatinib for 96 hours. Cell viability was assayed as described in Methods. Data are expressed as a percentage of the value for cells treated with vehicle control and are means of triplicates. The experiments were performed 3 times with similar results. (C) PC9 cells were treated with and without 500 nM afatinib for 60 minutes. *YES1* clones were maintained in 500 nM afatinib or grown without afatinib for 72 hours. Cell lysates were subjected to immunoblot analysis with antibodies against the indicated proteins.

**Fig. S3.**
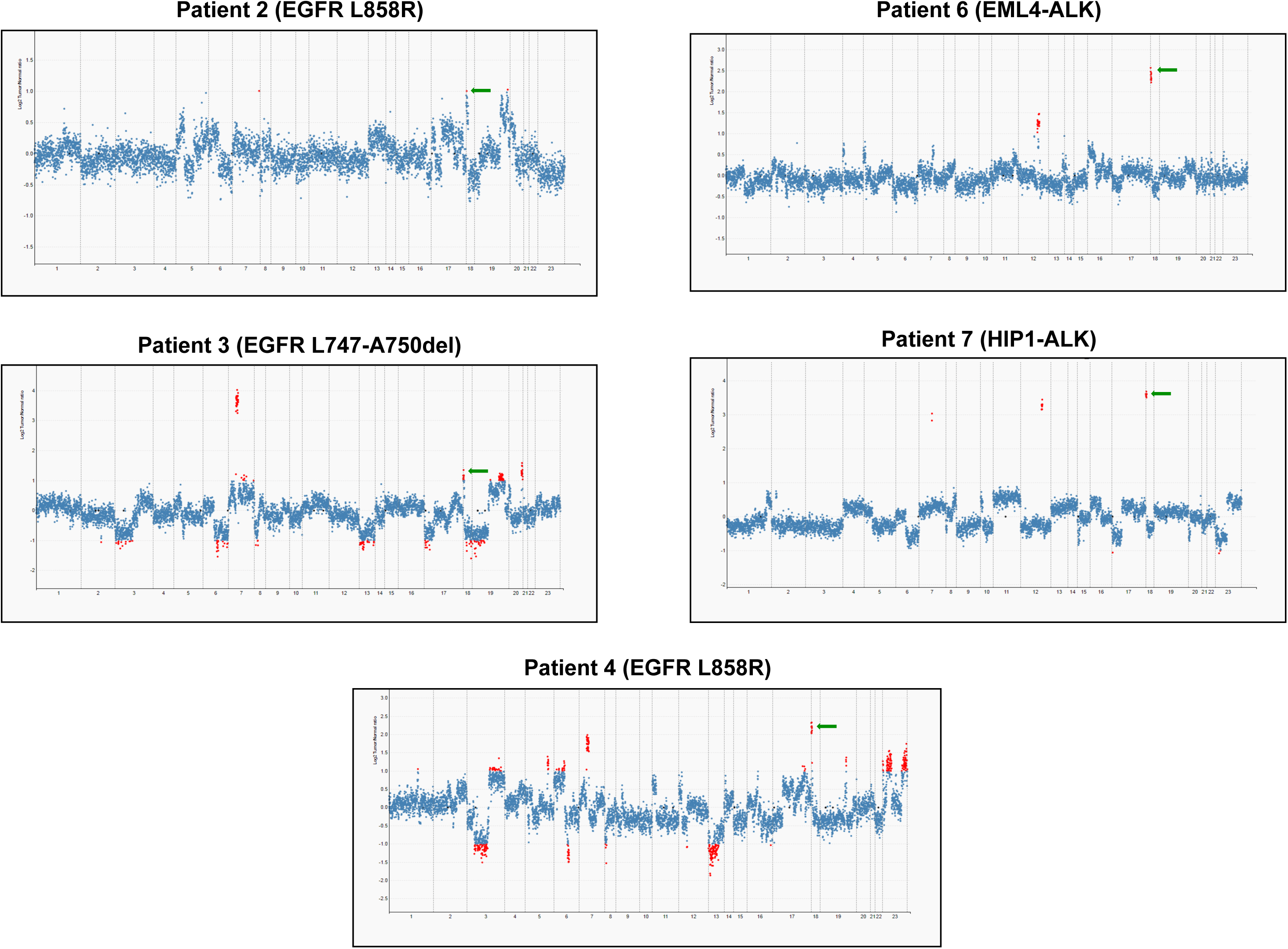
Amplification of *YES1* in post-TKI tumor samples from patients with acquired resistance to EGFR and ALK inhibitors. Copy number plots for post-TKI samples from patients 2, 3, 4, 6 and 7. Each dot represents a target region in the MSK-IMPACT targeted capture assay. Red dots are target regions exceeding a fold change cutoff of 2-fold. The log-ratios (y-axis) comparing tumor versus normal coverage values are calculated across all targeted regions (x-axis). Green arrows indicate focal amplification of *YES1* (11 coding exons targeted).

**Fig. S4.**
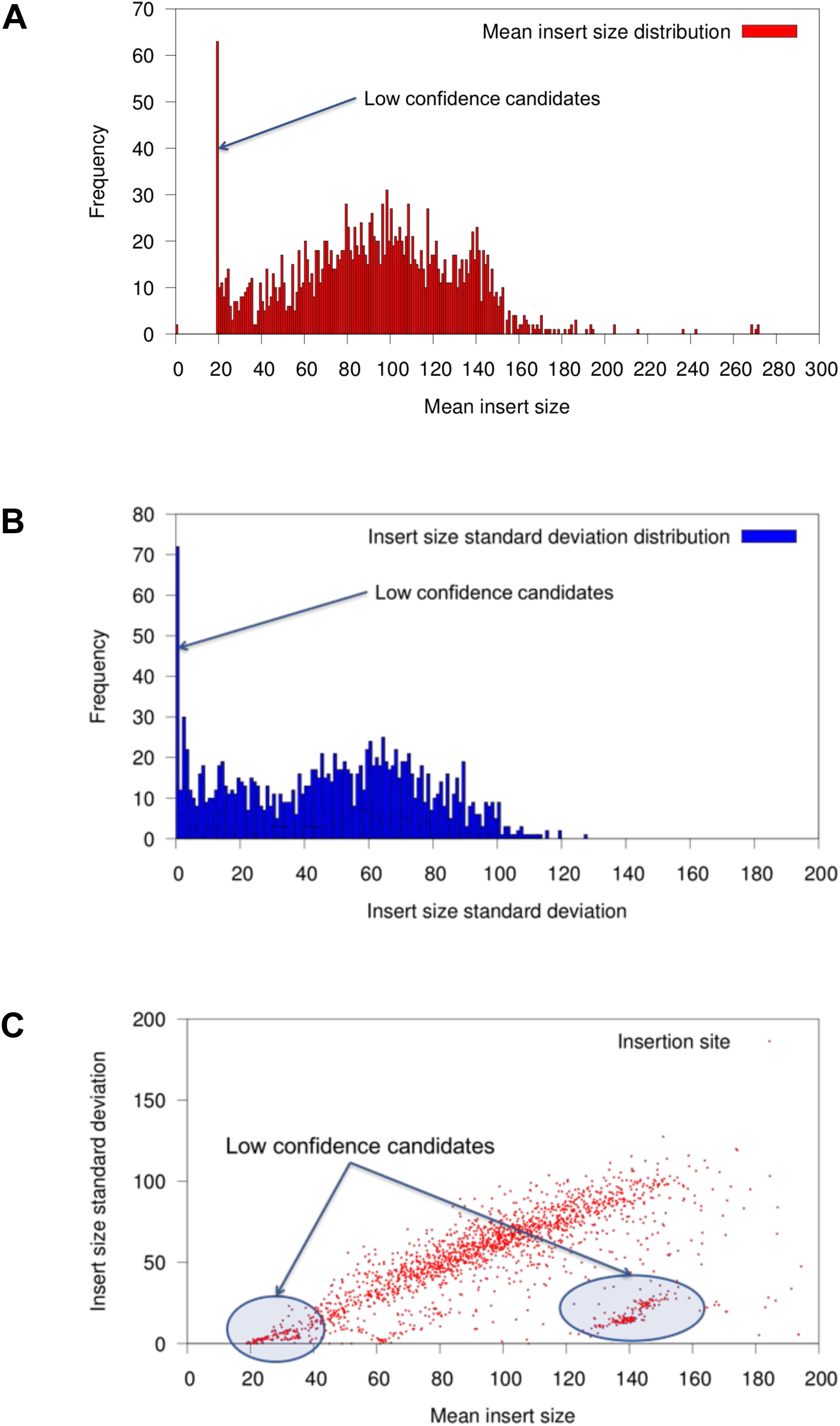
Mean insert size and standard deviation of insert size for candidate insertion sites. (A) Mean insert size distribution of candidate insertion sites. (B) Standard deviation of insert sizes for candidate insertion sites. (C) Correlation between mean insert size and standard deviation of insert size for the candidate insertion sites.

## Footnotes

1 Since there is no *YES2* gene and no other SFK gene name contains numerals, the authors suggested to the HUGO Gene Nomenclature Committee that the gene name be changed from *YES1* to *YES*. The committee did not agree to this change, noting that the use of “yes” in literature searches recovers numerous unrelated items. Regardless of the arguments for and against either *YES1* or *YES* as a gene name, the continued use of both *YES1* and *YES* within the scientific community necessitates the inclusion of both terms in literature searches to ensure retrieval of all publications that are relevant to the gene.

2 http://broadinstitute.github.io/picard/

3 https://github.com/pezmaster31/bamtools

## References

1. Kobayashi S, et al. (2005) EGFR mutation and resistance of non-smallcell lung cancer to gefitinib. N Engl J Med 352(8):786–792.

2. Pao W, et al. (2005) Acquired resistance of lung adenocarcinomas to gefitinib or erlotinib is associated with a second mutation in the EGFR kinase domain. PLoS Med 2(3):e73.

3. Wu SG, et al. (2016) The mechanism of acquired resistance to irreversible EGFR tyrosine kinase inhibitor-afatinib in lung adenocarcinoma patients. Oncotarget 7(11):12404–12413.

4. Campo M, et al. (2016) Acquired Resistance to First-Line Afatinib and the Challenges of Prearranged Progression Biopsies. J Thorac Oncol 11(11):2022–2026.

5. Thress KS, et al. (2015) Acquired EGFR C797S mutation mediates resistance to AZD9291 in non-small cell lung cancer harboring EGFR T790M. Nat Med 21(6):560–562.

6. Yu HA, et al. (2015) Acquired Resistance of EGFR-Mutant Lung Cancer to a T790M-Specific EGFR Inhibitor: Emergence of a Third Mutation (C797S) in the EGFR Tyrosine Kinase Domain. JAMA Oncol 1(7):982–984.

7. Sequist LV, et al. (2011) Genotypic and histological evolution of lung cancers acquiring resistance to EGFR inhibitors. Sci Transl Med 3(75):75ra26.

8. Yu HA, et al. (2013) Analysis of tumor specimens at the time of acquired resistance to EGFR-TKI therapy in 155 patients with EGFR-mutant lung cancers. Clin Cancer Res 19(8):2240–2247.

9. Engelman JA, et al. (2007) MET amplification leads to gefitinib resistance in lung cancer by activating ERBB3 signaling. Science 316(5827):1039–1043.

10. Zhang Z, et al. (2012) Activation of the AXL kinase causes resistance to EGFR-targeted therapy in lung cancer. Nat Genet 44(8):852–860.

11. Engelman JA, et al. (2006) Allelic dilution obscures detection of a biologically significant resistance mutation in EGFR-amplified lung cancer. J Clin Invest 116(10):2695–2706.

12. Ogino A, et al. (2007) Emergence of epidermal growth factor receptor T790M mutation during chronic exposure to gefitinib in a non small cell lung cancer cell line. Cancer Res 67(16):7807–7814.

13. Chmielecki J, et al. (2011) Optimization of dosing for EGFR-mutant nonsmall cell lung cancer with evolutionary cancer modeling. Sci Transl Med 3(90):90ra59.

14. Astsaturov I, et al. (2010) Synthetic lethal screen of an EGFR-centered network to improve targeted therapies. Sci Signal 3(140):ra67.

15. de Bruin EC, et al. (2014) Reduced NF1 expression confers resistance to EGFR inhibition in lung cancer. Cancer Discov 4(5):606–619.

16. Liao S, et al. (2017) A genetic interaction analysis identifies cancer drivers that modify EGFR dependency. Genes Dev 31(2):184–196.

17. Bivona TG, et al. (2011) FAS and NF-kappaB signalling modulate dependence of lung cancers on mutant EGFR. Nature 471(7339):523–526.

18. Ichihara E, et al. (2017) SFK/FAK Signaling Attenuates Osimertinib Efficacy in Both Drug-Sensitive and Drug-Resistant Models of EGFR-Mutant Lung Cancer. Cancer Res 77(11):2990–3000.

19. Sharifnia T, et al. (2014) Genetic modifiers of EGFR dependence in nonsmall cell lung cancer. Proc Natl Acad Sci U S A 111(52):18661–18666.

20. Krall EB, et al. (2017) KEAP1 loss modulates sensitivity to kinase targeted therapy in lung cancer. Elife 6.

21. DeNicola GM, Karreth FA, Adams DJ, & Wong CC (2015) The utility of transposon mutagenesis for cancer studies in the era of genome editing. Genome Biol 16:229.

22. Chen L, et al. (2013) Transposon activation mutagenesis as a screening tool for identifying resistance to cancer therapeutics. BMC Cancer 13:93.

23. Pandzic T, et al. (2016) Transposon Mutagenesis Reveals Fludarabine Resistance Mechanisms in Chronic Lymphocytic Leukemia. Clin Cancer Res 22(24):6217–6227.

24. Pettitt SJ, et al. (2013) A genetic screen using the PiggyBac transposon in haploid cells identifies Parp1 as a mediator of olaparib toxicity. PLoS One 8(4):e61520.

25. Chapeau EA, et al. (2017) Resistance mechanisms to TP53-MDM2 inhibition identified by in vivo piggyBac transposon mutagenesis screen in an Arf-/-mouse model. Proc Natl Acad Sci U S A 114(12):3151–3156.

26. Perna D, et al. (2015) BRAF inhibitor resistance mediated by the AKT pathway in an oncogenic BRAF mouse melanoma model. Proc Natl Acad Sci U S A 112(6):E536–545.

27. Choi J, et al. (2014) Identification of PLX4032-resistance mechanisms and implications for novel RAF inhibitors. Pigment Cell Melanoma Res 27(2):253–262.

28. Yusa K, Zhou L, Li MA, Bradley A, & Craig NL (2011) A hyperactive piggyBac transposase for mammalian applications. Proc Natl Acad Sci U S A 108(4):1531–1536.

29. Langridge GC, et al. (2009) Simultaneous assay of every Salmonella Typhi gene using one million transposon mutants. Genome Res 19(12):2308–2316.

30. Campbell JD, et al. (2016) Distinct patterns of somatic genome alterations in lung adenocarcinomas and squamous cell carcinomas. Nat Genet 48(6):607–616.

31. Takezawa K, et al. (2012) HER2 amplification: a potential mechanism of acquired resistance to EGFR inhibition in EGFR-mutant lung cancers that lack the second-site EGFRT790M mutation. Cancer Discov 2(10):922–933.

32. Li D, et al. (2008) BIBW2992, an irreversible EGFR/HER2 inhibitor highly effective in preclinical lung cancer models. Oncogene 27(34):4702–4711.

33. Soria JC, et al. (2018) Osimertinib in Untreated EGFR-Mutated Advanced Non-Small-Cell Lung Cancer. N Engl J Med 378(2):113–125.

34. Yu HA, et al. (2018) Concurrent alterations in EGFR-mutant lung cancers associated with resistance to EGFR kinase inhibitors and characterization of MTOR as a mediator of resistance. Clin Cancer Res (in press).

35. Ohashi K, et al. (2012) Lung cancers with acquired resistance to EGFR inhibitors occasionally harbor BRAF gene mutations but lack mutations in KRAS, NRAS, or MEK1. Proc Natl Acad Sci U S A 109(31):E2127–2133.

36. Suda K, et al. (2010) Reciprocal and complementary role of MET amplification and EGFR T790M mutation in acquired resistance to kinase inhibitors in lung cancer. Clin Cancer Res 16(22):5489–5498.

37. Takeda T, et al. (2017) Yes1 signaling mediates the resistance to Trastuzumab/Lap atinib in breast cancer. PLoS One 12(2):e0171356.

38. Johnson ML, et al. (2011) Phase II trial of dasatinib for patients with acquired resistance to treatment with the epidermal growth factor receptor tyrosine kinase inhibitors erlotinib or gefitinib. J Thorac Oncol 6(6):1128–1131.

39. Crystal AS, et al. (2014) Patient-derived models of acquired resistance can identify effective drug combinations for cancer. Science 346(6216):1480–1486.

40. Cheng DT, et al. (2015) Memorial Sloan Kettering-Integrated Mutation Profiling of Actionable Cancer Targets (MSK-IMPACT): A Hybridization Capture-Based Next-Generation Sequencing Clinical Assay for Solid Tumor Molecular Oncology. J Mol Diagn 17(3):251–264.

41. Ross DS, et al. (2017) Next-Generation Assessment of Human Epidermal Growth Factor Receptor 2 (ERBB2) Amplification Status: Clinical Validation in the Context of a Hybrid Capture-Based, Comprehensive Solid Tumor Genomic Profiling Assay. J Mol Diagn 19(2):244–254.

42. Shen R & Seshan VE (2016) FACETS: allele-specific copy number and clonal heterogeneity analysis tool for high-throughput DNA sequencing. Nucleic Acids Res 44(16):e131.

43. Rizvi H, et al. (2018) Molecular Determinants of Response to Anti-Programmed Cell Death (PD)-1 and Anti-Programmed Death-Ligand 1 (PD-L1) Blockade in Patients With Non-Small-Cell Lung Cancer Profiled With Targeted Next-Generation Sequencing. J Clin Oncol 36(7):633–641.

## References

1. Li H. (2013) Aligning sequence reads, clone sequences and assembly contigs with BWA-MEM. 1303.3997v1 [q-bio.GN]

